# FOXO regulates neuromuscular junction homeostasis during *Drosophila* aging

**DOI:** 10.1101/2020.04.13.040121

**Authors:** Allison Birnbaum, Kai Chang, Hua Bai

## Abstract

The transcription factor FOXO is a known regulator of lifespan extension and tissue homeostasis. It has been linked to the maintenance of neuronal processes across many species, and has been shown to promote youthful characteristics by regulating cytoskeletal flexibility and synaptic plasticity at the neuromuscular junction (NMJ). However, the role of FOXO in aging neuromuscular junction function has yet to be determined. We profiled adult Drosophila FOXO-null mutant abdominal ventral longitudinal muscles and found that young mutants exhibited morphological profiles similar to those of aged wild-type flies, such as larger bouton areas and shorter terminal branches. We also observed changes to the axonal cytoskeleton and an accumulation of late endosomes in FOXO null mutants and motor neuron-specific FOXO knockdown flies, similar to those of aged wild-types. Motor neuron-specific overexpression of FOXO can delay age-dependent changes to NMJ morphology, suggesting FOXO is responsible for maintaining NMJ integrity during aging. Through genetic screening, we identify several downstream factors mediated through FOXO-regulated NMJ homeostasis, including genes involved in the p38-MAPK pathway. Interestingly, the phosphorylation of p38 and ERK were increased in the motor neuron-specific FOXO knockdown flies, suggesting FOXO acts as a suppressor of MAPK activation. Our work reveals that FOXO is a key regulator for NMJ homeostasis, and it maintains NMJ integrity by repressing MAPK signaling during aging.

## Introduction

Aging involves the progressive functional decline of cellular mechanisms and tissue integrity (Rose 1994; Partridge and Barton 1996; Lopez-Otin et al 2013) In the adult brain, this results in a gradual decline of synaptic contacts to skeletal muscle tissue, resulting in a loss of strength and muscle mass (Hall and Sanes 1993). The neuromuscular junction (NMJ) serves as the synaptic interface between the branched terminals of motor neurons and the skeletal muscle fibers (Punga and Ruegg 2012). The NMJ is highly dynamic in response to cellular signals and stressors, and a dysregulation of molecular processes in both the pre- and post-synaptic regions can lead to the onset of neurodegenerative diseases (Gonzalez-Freire et al 2014; Monani and De Vivo 2014). With aging there is impairment of regulatory systems such as autophagy and redox homeostasis at the NMJ, which can lead to reactive oxygen species (ROS) accumulation followed by organelle damage and cell death (Li et al 2018; Stefanatos and Sanz 2018). Synaptic plasticity, which allows for the maintenance of functional activity to protect against degeneration, can decline during aging, resulting in decreased neuronal responsiveness and synaptic deterioration (Bergado and Anlmaguer 2002; Kempsell and Fieber 2015; Wagner et al 2015).

The *Drosophila* NMJ synapse utilizes glutamate as the primary neurotransmitter, and has postsynaptic densities that behave similarly to the AMPA-type receptors in the vertebrate central nervous system (Menon et al 2013). The Drosophila neuromuscular junction system has been well stereotyped in larvae, but has only more recently begun to be characterized in the adult organism (Rivlin et al 2004; Hebbar et al 2006; Beramendi et al 2008; Wagner et al 2015; Lopez-Arias et al 2017). These studies have also discovered a number of morphological changes to the NMJ that occur under normal aging conditions, and can aid in understanding the mechanisms involved in age-dependent neuronal impairment.

The FOXO transcription factor family plays an important role in metabolism and development as well as stress resistance and lifespan (Accili and Arden 2004; Greer and Brunet 2005). FOXO has long been implemented as a “longevity gene” and as a modulator of protein homeostasis (Kenyon et al 1993, Martins et al 2016). FOXOs can also function as regulators of neuronal homeostasis (Paik et al 2009; Kim and Webb 2017; Schaffner et al 2018). FOXO-mediated pathways permit neuronal plasticity, which have downstream implications on cellular behavior and neuronal morphology (McLaughlin and Broihier 2018). *Drosophila* FOXO (dFOXO, hereafter FOXO) promotes cytoskeletal dynamics at the neuromuscular junction during development, where it responds to stimulation to allow rapid structural reorganization. In the Drosophila larvae NMJ, loss of FOXO results in increased stability of microtubules through the upregulation of kinesin motor protein regulators that also control axon growth (Nechipurenko and Broihier 2012; McLaughlin et al 2016). In mammals, certain FOXO isoforms show transcriptional regulation of cytoskeletal polarity genes during development (de la Torre-Ubieta et al 2010). With aging, FOXOs are also implicated in regulating autophagy in neurons under Parkinson’s disease models (Xu et al 2011; Pino et al 2014; Schaffner et al 2018). It has also been shown that aging results in the activation of pro-inflammatory signals and proteotoxic stress in mice brains, and the depletion of neuronal FOXOs causes premature occurrences of these events and induces motor function impairment (Hwang et al 2018). Additionally, loss of FOXO impacts synaptic function by delaying neurotransmitter release in both larval and adult drosophila neuromuscular junctions (Howlett et al 2008; Mahoney et al 2016). Aged synapses also show a reduction in evoked response and neurotransmission (Segal 1982; Porras and Mora 1995; Liu et al 2013). These results indicate that FOXO is an important regulator of neuronal function, and that it has influence on functional and structural integrity in regulating the connection between the brain and the muscle tissue.

Our current knowledge of FOXO supports the hypothesis that FOXO activity promotes youthful NMJ morphology and provides necessary regulation of pathways involved in maintaining synaptic integrity and plasticity. However, how FOXO impacts functional regulation of the NMJ during aging still remains to be elucidated. Here, we show that loss of FOXO causes morphological alterations to the NMJ in young adults, and shares phenotypic characteristics with those of aging control organisms. We find that knockdown of FOXO specifically in the motor neuron results in changes to microtubule morphology and an upregulation of late endocytic vesicles. These events are also observed in middle-aged flies, indicating loss of FOXO may lead to a disruption of motor neuron homeostasis. Overexpression of FOXO delays age-related changes to NMJ morphology. Additionally, we identified potential FOXO downstream pathways that may promote aging and decline in synaptic function in the adult motor neurons. These findings provide insight into the role FOXO plays in maintaining morphological and synaptic plasticity at the aging neuromuscular junction.

## Results

### Loss of FOXO causes morphological changes to bouton size and branch length that correspond to aging

Normal aging causes alterations to the morphology of boutons and branches at the adult Drosophila neuromuscular junction (Beramendi et al 2007, Wagner et al 2015, Lopez-Arias et al 2017). Based on previous literature, we selected three time points post eclosion, 5 days, 25 days, and 40 days to represent young, middle aged, and old adult flies. We visualized the A3 abdominal ventral longitudinal muscle (VLM) segment using immunofluorescence staining. We stained flayed adult abdominal pelts with a monoclonal antibody against bruchpilot (BRP) to mark active zones (AZs) and an antibody against horseradish peroxidase (HRP) to visualize neuronal tissue (Figure 1A). To obtain overall bouton number, we used an antibody against Disc-large (Dlg) to mark post-synaptic regions. Using the two NMJ’s present at each A3, we quantified the six metrics characterized by Wagner et al 2015 and found a significant difference between ages for several metrics, as expected. A significant decrease in branch length was observed with aging, along with a decrease in bouton number(Figure 1B, Table 1). We also found an expansion of bouton area with age, and this corresponded to an increase in number of AZs per bouton(Figure 1C-D, Table 1). However, the number of AZs observed in a 100um^2^ region did not change between ages, which has been shown to remain constant during a normal progression of aging (Table 1, Wagner et al 2015).

**Table 1.**
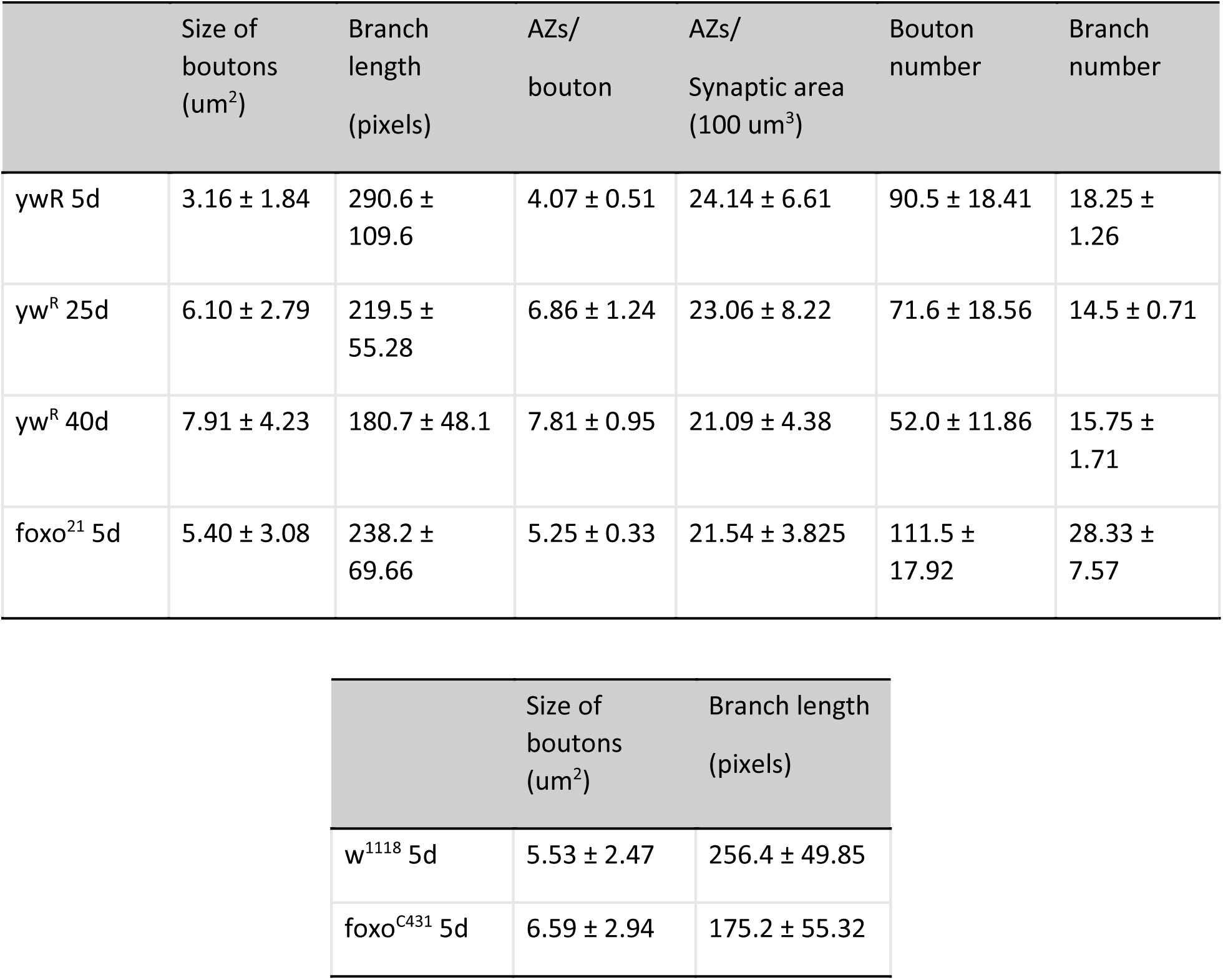
Quantification of NMJ morphological characteristics of aging. yw^R^ is the control line for foxo^21^. Average bouton number: 5-day yw^R^ v 5-day foxo^21^ (n.s.). 5-day yw^R^ v 25-day yw^R^ (n.s.). 5-day yw^R^ v 40-day yw^R^ (P< 0.05). 25-day yw^R^ v 5-day foxo^21^ (P< 0.05). Branch number: 5-day yw^R^ v 5-day foxo^21^ (P< 0.05). 5-day yw^R^ v 25-day yw^R^ (n.s.). 5-day yw^R^ v 40-day yw^R^ (n.s.). Bouton number and branch number represent both NMJs present at the A3 abdominal segment.

**Figure 1.**
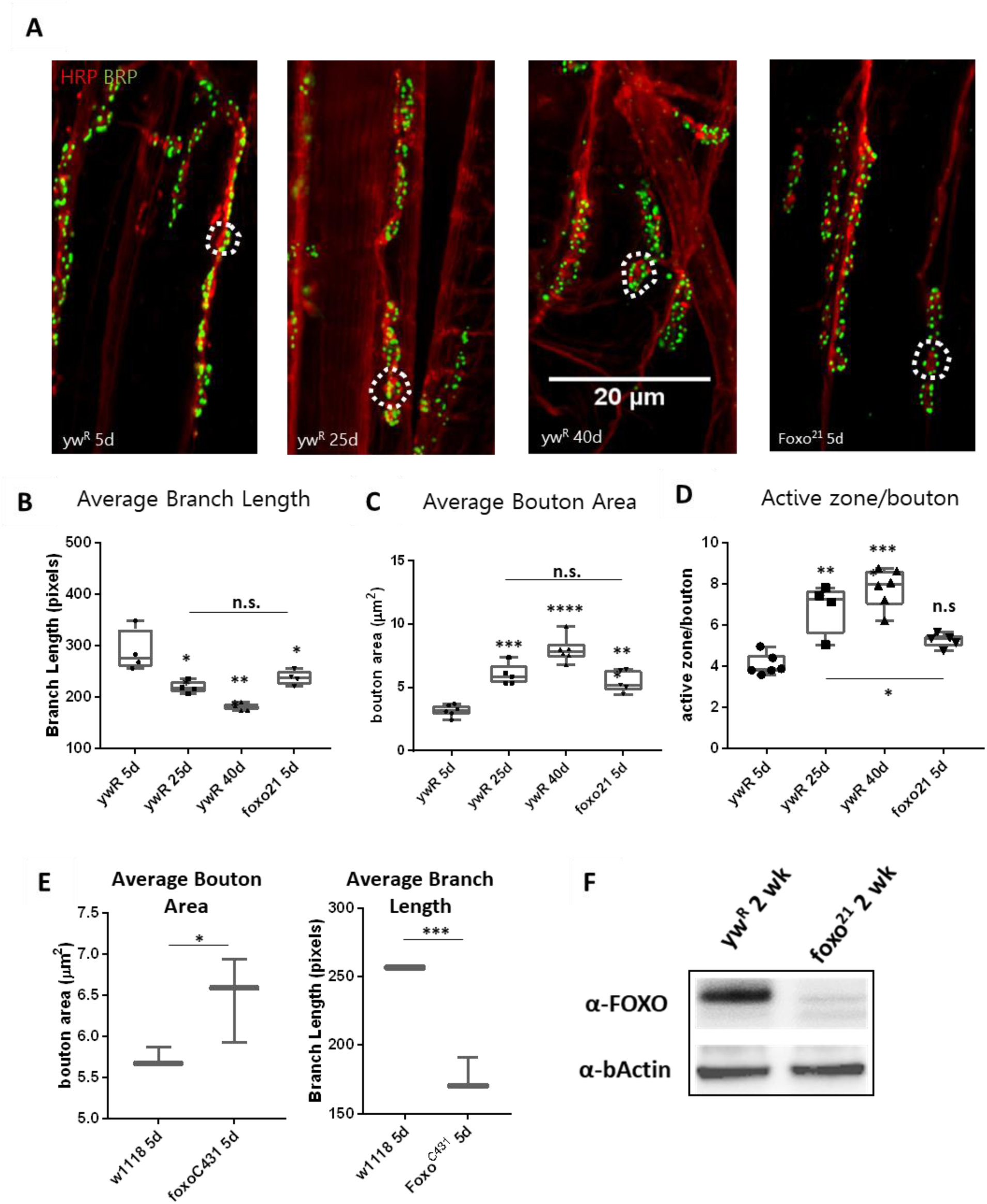
Loss of FOXO results in middle-aged NMJ morphology in the abdominal ventral muscle. **A.** Representative images of the Abdominal VLM A3 segment for axon terminals with indicated genotypes and ages colabeled with Anti-BRP and anti-HRP. Bar 20 μM. **B.** Average branch length. *P<0.05, ** P<0.01. Bouton area for 5-day *foxo* flies is significantly larger than 5-day controls, but is comparable to 25-day control flies. ** P<0.01. **C.** Average bouton area ** P<0.01, ***P>0.001, ****P< 0.0001. **D.** Quantification of changes in active zone number per bouton. *P<0.05, ** P<0.01, ***P<0.001. **E.** Quantification of bouton area and branch length for 5-day old w1118 control flies and foxo^c431^ CRISPR FOXO-null flies. *P<0.05, ** P<0.01. **F.** Western analysis of FOXO protein abundance in control and *Foxo*^*21*^ mutant whole-body tissue. Number of animals analyzed for NMJ quantification foxo^21^ 5d n=5, ywR 5d n=6, 25d n=5, 40d n=6. number of boutons quantified per sample analyzed foxo^21^ 5d n=100, ywR 5d n=177, 25d n=75, 40d n=96. number of branches analyzed foxo^21^ 5d n=39, ywR 5d n=39, 25d n=39, 40d n=40

We next examined the effect loss of FOXO had on NMJ morphology in the adult. We used the strong hypomorph *foxo^21^* allele and dissected the VLM of 5-day post eclosion animals and characterized the A3 muscle as described previously (Figure 1A). We found that *foxo* mutants had significantly larger bouton areas and shorter branches than their 5-day wild-type counterparts. However, the bouton area and branch length of *foxo* mutants was not significantly different from those seen in middle age control flies (Figure 1B-C, Table 1). The average active zone number between young wild-type and *foxo* mutants was not significantly different, but the *foxo*-null mutants had more average AZs per bouton compared to controls (Figure 1D, Table 1, Figure S1). When evaluating the raw total number of active zones per bouton, a significant difference was detected between 5-day controls and *foxo* mutants, but not between 25-day controls and 5-day *foxo* mutants (Figure S1). Additionally, we did not detect a significant difference in overall AZs per synaptic area in *foxo* mutants when compared to the three measured control age groups. We did find that *foxo* mutants on average had more branches and a higher bouton number than wild-type flies (Table 1), which may be a compensatory response to reduced neurotransmission under excitatory conditions seen in *foxo* mutants (Howlett et al 2008, Mahoney et al 2016, McLaughlin and Broihier 2017). We also examined bouton size and branch length in 5-day old flies containing a null *foxo^C431^* allele generated through CRISPR-Cas9 (Birnbaum et al 2019). We found these null mutants also had enlarged boutons and shorter branch lengths compared to controls at a young age (Figure 1E). We validated that foxo protein levels were indeed reduced in our mutants (Figure 1F). These results suggest foxo promotes aging morphologies of boutons and synaptic branches at the neuromuscular junction.

### Loss of FOXO results in altered cytoskeleton structure in the adult neuromuscular junction

Axonal degeneration is a normal consequence of aging, and arises from a number of alterations to cellular and molecular pathways (Salvadores et al 2017). This can result in a withdrawal of axons from their synaptic sites, caused by changes in the underlying cytoskeleton (Manini et al 2013, He et al 2002). Foxo proteins are known to regulate motor neuron microtubule dynamics during development across many species, and can cause a decrease in microtubule stability, allowing for increased plasticity (de la Torre-Ubieta et al 2010, Nechipurenko and Broiheir 2012, McLaughlin et al 2016). However, how foxo effects microtubule dynamics in a post-developmental model has not been examined. We used an antibody against acetylated alpha tubulin (Ac-tub) which marks stable microtubules at our three aging time points. We constructed a 3D model of each A3 segment and assigned all branches containing acetylated a-tubulin one of two characteristics; straight, or undulating. We quantified the percentage of each tubulin morphology found the undulating morphology significantly increases between 5 and 25 days, and between 25 and 40 days (Figure 2A, Table 2). In examining the wild-type flies, we found at the young time point around 70% of branches had a straight tubulin structure. As wild-type flies aged, we saw an increase in sinusoidal bends persisting throughout the branches, creating an undulating pattern (Figure S2). This morphology can be associated with axonal retraction (He et al 2002). These results show that there are changes that occur to the axonal microtubule structure with aging.

**Table 2.** Quantification of acetylated alpha-tubulin structure. Significance values are compared to young control for each grouping

|  | Straight | Undulating |
| --- | --- | --- |
| yw<sup>R</sup> 5d | 69.24% | 30.76% |
| yw<sup>R</sup> 25d | 40.38%* | 59.62%* |
| yw<sup>R</sup> 40d | 30.00%** | 70.00%** |
| foxo<sup>21</sup> 5d | 46.83%* | 53.17%* |
| ok6 > control1 wk | 67% | 33% |
| ok6 > foxo-RNAi 1wk | 34.27%*** | 65.73%*** |
| ok6 > control 25d | 38.64%** | 61.36%** |

**Figure 2.**
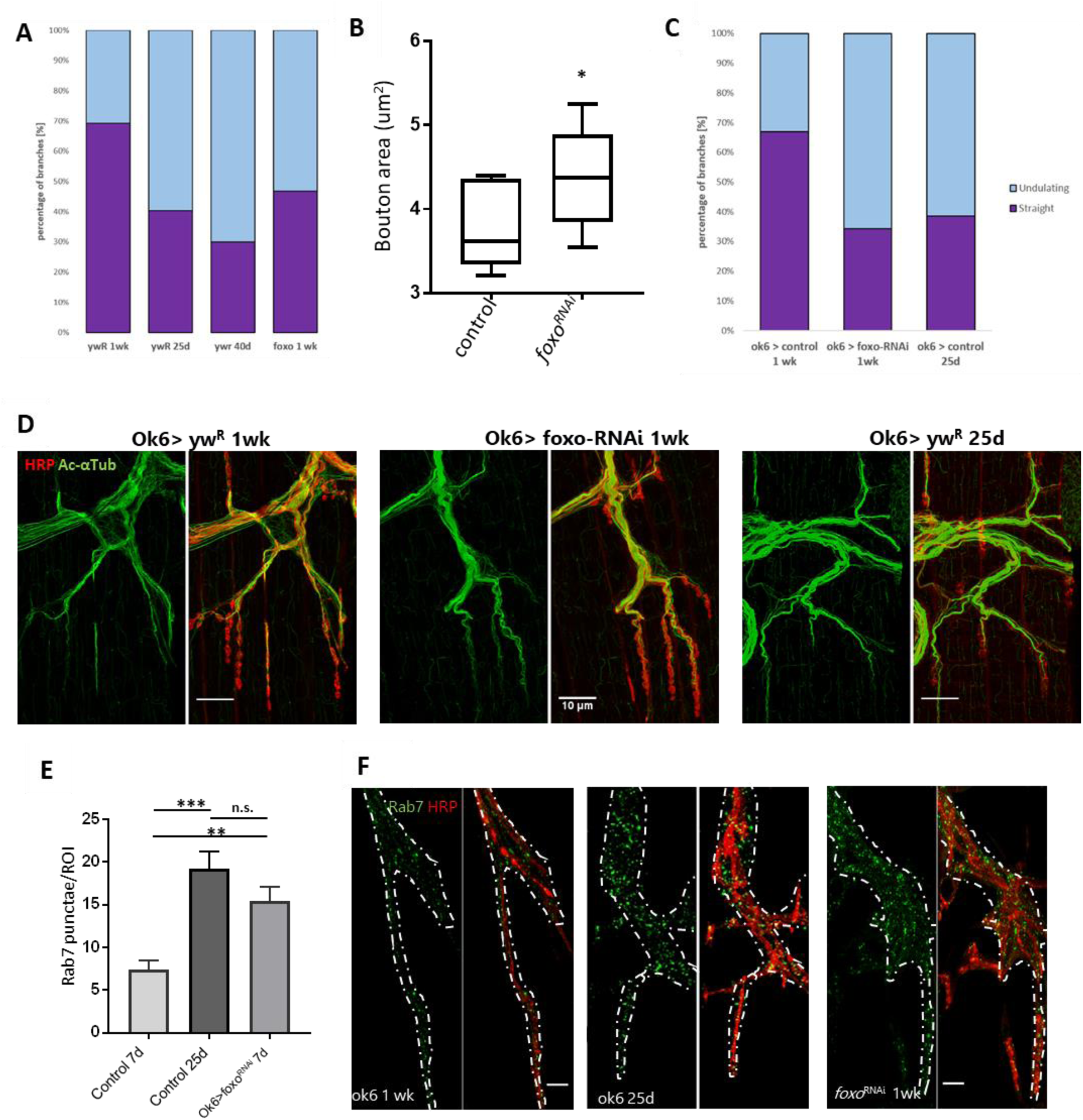
Aging and loss of FOXO cause changes in microtubule structure and endocytic activity. **A.** Quantification of anti-Ac-Tub at A3 VLM at control aging timepoints and in *foxo* mutant. The mean fraction of straight branches was significantly reduced from the young control for all other columns (P< 0.05). **B.** Average bouton area for ok6-GAL4>yw^R^ compared to ok6-GAL4>foxo-RNAi #1 (*P< 0.05). **C.** Quantification of anti-Ac-Tub at A3 VLM at specified genotypes and ages. The % of straight branches of all other columns was significantly reduced from the young control (P< 0.05). **D.** Representative images of axon terminals with indicated genotypes and ages. Colabeled with Anti-Ac-Tub and anti-HRP. Bar 10 μM. **E.** Number of Rab7 punctae detected per ROI in A3 segment. **P< 0.01, ***P< 0.001, n.s.-not significant. **F.** Representative images of main axon branchpoint with indicated genotypes and ages. Colabeled with Anti-Rab7 and anti-HRP. Bar 10 μM.

We next examined our young adult *foxo* mutants for any disruptions to the axonal microtubule morphology. *Foxo* mutants exhibited an increased number of undulating microtubules within branches, and had significantly fewer straight microtubule bundles than the control at 5 days (Figure S2). When the *foxo* mutants were compared to the 25-day controls, there was no significant difference detected between the percentage of both undulating and straight microtubule branches (Figure 2A). The *foxo^21^* line contains a whole body hypomorph, and we could not determine if this change in microtubule morphology was a result of motor neuron FOXO activity. Therefore, we used a motor neuron specific *Ok6-Gal4* (gift from E. McNeill Lab) to knock down *foxo* specifically in the motor neuron. We quantified bouton size between control and foxo-knockdown flies and found that knock-down of foxo in the motor neuron induces an increase in bouton size (figure 2B). We stained A3 VLMs against Ac-tub, and at one-week post eclosion, and observed nearly a two-fold decrease in the number of straight branches in knockdown flies compared to controls (*Ok6-Gal4>ywR)* (figure 2C, table 2). We also compared our knockdown to 25-day controls and found there to be no significant difference between the percentage of undulating and straight microtubules within axonal branches (figure 2C-D, table 2). Through knocking down foxo in the motor neuron, we have uncovered that it is foxo motor neuron activity that influences bouton size and axonal microtubule morphology, and that this early undulating pattern observed in foxo mutants suggest it is involved in maintaining youthful characteristics of the neuronal cytoskeleton.

### Loss of foxo causes impairment of the endocytic pathway in the adult neuromuscular junction

Aging is also known to cause disruption of endosome-lysosome trafficking which can result in neurodegeneration, and increases in the number of early endosomes and multivesicular bodies (prerequisites for late endosomes) at the synapse (Boaro et al 1998, Nixon et al 2008, Wagner et al 2015, Colacurcipo and Nixon 2016). In *C. elegans*, late endosome formation is suppressed in organisms with delayed aging (Richardson et al 2019). Additionally, late endosomes undergo retrograde transport to degrade cargo, and this transport declines with age (Milde et al 2015, De Vos and Hafezparast 2017). To observe changes in the endocytic degradation pathway, we used an antibody against rab7 to mark late endosomes. We performed immunofluorescent staining at the A3 VLM on 5 day and 25-day adults. To our surprise, we observed rab7-marked punctae around the axon branchpoint site, and noticed a visible increase at this site in middle-aged flies. To quantify the number of rab7 vesicles, we generated circular regions of interest (ROIs) of the same size and used HRP to identify z-slices containing the axon. The slice with the highest punctae number for each ROI was collected and averaged for each specimen. We found the 25-day adult flies had more than 2-fold more Rab7 punctae per ROI associated with the axon branch than the one-week adults, demonstrating there is an increase in the number of late endosomes during aging (Figure 2E-F).

We next examined our ok6-gal4 foxo knockdown flies for alterations in late endosome signaling. We used the same method, we quantified neuronal Rab7 punctae. We saw foxo KD flies also had significantly more punctae per ROI than the young controls (Figure 2 G, H). We tested a second FOXO-RNAi line (FOXO-RNAi #2) and again observed significantly more rab7 punctae at the branchpoint than controls (Figure S3). Overall, both aging and foxo knockdown result in an increase in the number of late endosomes around the branchpoint of motor neuron, suggesting some cellular disruption in both instances.

### Overexpression of FOXO rescues NMJ aging phenotypes

To test if FOXO is the cause of this increase in late endosome number, we overexpressed foxo using a UAS system. This was driven by *Ok6-GAL4* to ensure expression specifically in the motor neuron. At one-week post eclosion, no differences were detected in rab7 number between controls and overexpressing foxo flies (Figure 3 A,B,C). However, at 25 days, controls had accumulated significantly more late endosomes along the axon compared to overexpression lines (Figure A’ B’, C). We next examined how over expression of foxo impacts bouton morphology with aging, and found controls to have greater average bouton size compared to foxo overexpression at 25 days (Figure 3D). To test for the effectiveness of foxo overexpression, we used a daughterless inducible gene switch driver line. To activate the gene switch, we fed flies 200mM of Mifepristone (RU486), and used the whole fly body for RNA extraction. We examined expression of the known foxo target Thor (4EBP), and found a 2-fold increase in expression in RU486 fed flies, indicating this line can sufficiently induce foxo overexpression (Figure S3). Altogether, these results indicate that foxo expression can be beneficial to preserve motor neuron morphology and maintain neuronal endocytosis.

**Figure 3.**
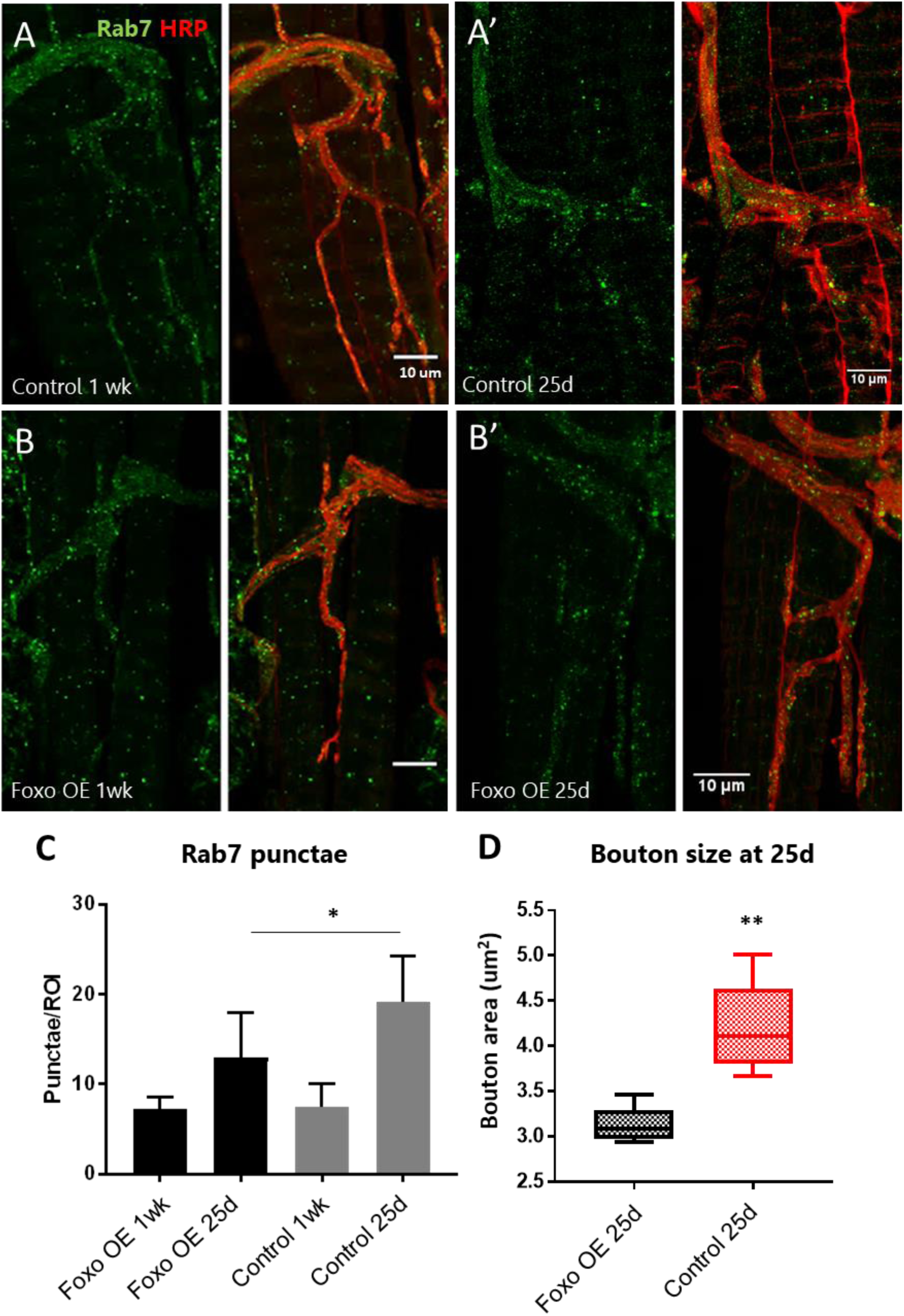
Overexpression of motor neuron FOXO delays aging phenotypes. **A-B.** Representative figures of control and FOXO-overexpression at 1 week, (A’-B’) and 25 days. Colabelling with anti-Rab7 and anti-HRP. Scale bar 10μm. **C.** Rab7 quantification for A3 ROIs for indicated genotypes and ages *P < 0.05. **D.** Average bouton area for 25-day old FOXO overexpression and control **P<0.01.

### MAPK and Activin signaling can rescue knockdown induced NMJ disruptions

In order to understand how loss of FOXO disrupts NMJ homeostasis, we sought to uncover the pathway that is downstream of FOXO regulation in the motor neuron. To accomplish this, we utilized a combination of genomic and genetic approaches to identify candidate genes. We surveyed foxo ChIP data and compared this with transcriptome data from head tissue of foxo-null mutants and aging controls to select candidates (Birnbaum et al 2019). Only foxo bound genes from our ChIP-seq that had a fold change 1.5 greater than the input baseline was used for comparison. For transcriptional data, we selected genes with a significant p-value and log2fold of

±0.25, indicating a 1.25 change in expression from both RNA-seq datasets. We identified 207 genes shared between all three data sets (Figure 4A, Supplementary Table). This gene set was enriched for biological processes involving neurotransmitter secretion and cytoskeletal organization. We also observed enhancement of the Phosphatidylinositol signaling pathway (Supplementary Table). We expanded our search to include genes that were bound by foxo and had altered transcription either with loss of foxo or with aging, generating a gene set list of 980 genes. We again found biological processes involving neurotransmission and axonal guidance, however we also found synaptic vesicle endocytosis and Rho signaling to be enriched among this gene set as well. Kegg pathways highly represented among this data include MAPK, WNT, Hippo, and TGF-b signaling. When we examined the 439 genes that were shared between both transcriptome data sets, but were not present in the foxo ChIP analysis, we saw enrichment of oxidation-reduction processes, and immunity regulating pathways (Supplemental Table). These results suggest foxo may serve as an indirect regulator in these pathways, or may require specific activation to bind DNA for transcriptional control.

**Figure 4:**
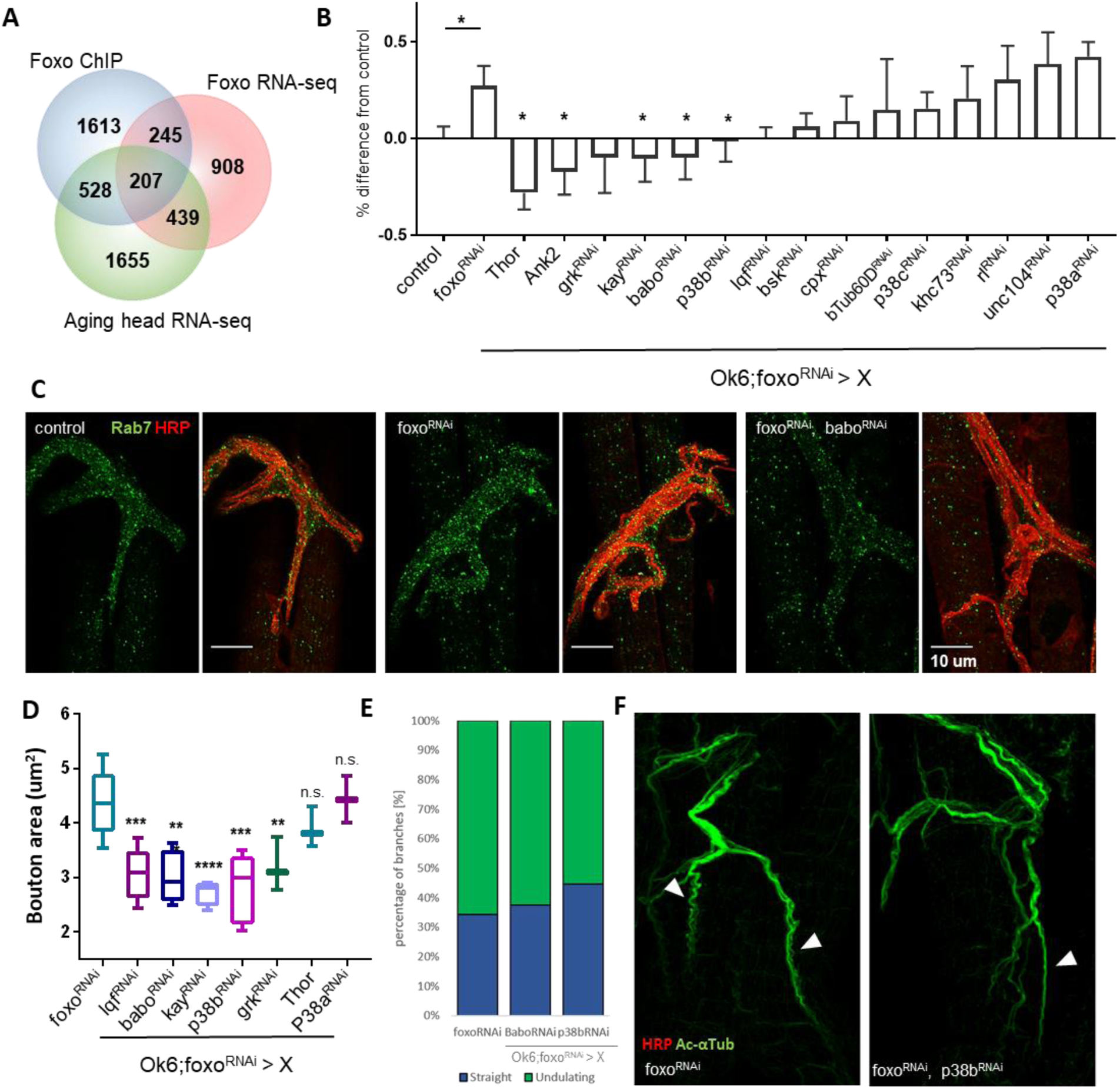
MAPK acts downstream of FOXO to regulate NMJ pathways. **A.** Venn diagram overlap of ChIP-seq, head foxo^C431^ RNA-seq, and aged head RNA-seq: For RNA-seq data {-0.15 log2FC > 0.15, P < 0.05}. {FOXO ChIP FC >1.5, FDR< 0.05}. **B.** Quantification of Rab7 for double mutant flies. Values are set as a percentage with ok6>yw^R^ as baseline. All significant values are compared to foxo^RNAi^ (in ok6-GAL4 background) *P<0.05. **C.** Representative images of genetic screening. Genotypes as indicated. Scale bar 10μm. **D.** Average bouton size quantification for indicated genotypes. Control in Ok6-GAL4 background. *P<0.05, **P<0.01, ***P< 0.001, ****P< 0.0001, n.s. – not significant. **E.** Quantification of the mean fraction of anti-Ac-Tub branch morphology. Mean fraction of straight branches is significantly higher in p38b^RNAi^. P<0.05. **F.** Representative figures of anti-Ac-Tub staining. Arrows indicate undulating (left figure) and straight (right figure) branches. Scale bar 10μm.

We generated a list of 15 candidates that we hypothesize to function downstream of foxo based on our genomic analysis, along with targets that have shown overlap between foxo activity and synaptic integrity throughout academic literature (Table 3). In our in foxo-null head transcriptome dataset, we observed upregulation of Activin and MAPK regulating genes. These pathways were also enriched in the foxo ChIP-seq dataset. In the follow-up genetic screening, we opted to use rab7 as a marker due to the ease of rapid quantification. We generated a fly line that containing the *Ok6-gal4* driver with a foxo-RNAi to perform a genetic screen to rescue the accumulation of rab7. Our generated fly showed a 27% increase in the number of late endosomes associated with the axon branchpoint compared to a wild-type (Figure 4B). For our assay, we created 13 double knockdown lines containing a knockdown of foxo as well as our candidate genes. We also used two overexpression lines with our ok6;FOXO knockdown. One was for Thor, which is a known FOXO target gene and has been shown to rescue *foxo* mutant neurotransmission defects (Junger et al 2003, Mahoney et al 2016). The other overexpression line we used was for the microtubule associated protein Ankyrin2 (Gift from Dubreuil lab, Weber et al 2019), which showed a decline in expression in foxo mutants and with aging in our transcriptome data, and was highly enriched in our ChIP-seq (Table 3). We used the same method from our previous results to quantify the number of late endocytic vesicles associated with the axon for each cross. Using a student t test, we identified four genes that resulted in a significant reduction in late endosome number compared to the foxo KD line. For our overexpression lines, we saw 28% and 17% less rab7 than the control from Thor and Ankryin2 respectively. We also found a significant reduction in rab7 marked vesicles when we expressed RNAi-mediated knockdown of the JNK transcription factor *Kay* (*Kayak*), and the TGF-b receptor *babo* (*baboon*). Both are known to have regulatory effects on axonal transport (Ellis et al 2010, Dreup and Nechiporuk 2013). We also saw a decline in the number of rab7 vesicles in p38b and grk double RNAi flies that were not significant, but were reduced near the baseline. Of the three p38 isoforms, only p38b was able to show this level of reduction. This isoform has also been implicated in axonal transport (Gibbs et al 2018). These results indicate that TGF-b and MAPK signaling are activated upon loss of foxo, and can disrupt normal endocytic activity associated with the motor neuron. After identifying potential rescues of late endosome accumulation in our genetic screen, we tested our positive hits for their ability to rescue other foxo KD induced phenotypes.

**Table 3:** Downstream candidate genes from FOXO ChIP-seq and RNA-seq

| Gene symbol | Gene name | function | ChIP FC | FOXO RNA-seq log2FC | Aging head RNA-seq log2FC |
| --- | --- | --- | --- | --- | --- |
| rl | rolled | ERK | 4.73 | 0.343 | 0.159 |
| Lqf | liquid facets | Epsin1 | 2.67 | 0.386 | 1.132 |
| Unc-104 | uncoordinated-104 | Kinesin/KIF1A | 2.33 | -0.388 | 0.327 |
| Khc-73 | Kinesin heavy chain 73 | Kinesin | 1.94 | -0.23 | 0.106 |
| Ank2 | Ankyrin 2 | Microtubule associated protein | 3.48 | -0.433 | 0.468 |
| Cpx | complexin | SNARE regulator | 3.68 | -0.22 | 1.001 |
| bTub60D | beta-Tubulin at 60D | Microtubule building block | 1.91 | 1.442 | -1.147 |
| Babo | baboon | Activin receptor | 3.71 | 0.244 | 0.258 |
| Grk | gurken | ERK ligand | 2.61 | 0.292 | -0.560 |
| Thor | Thor | 4EBP | 3.24 | -0.225 | -0.137 |
| Bsk | basket | JNK | - | -0.343 | 0.529 |
| Kay | kayak | JNK Transcription factor | 2.55 | -0.059 | -0.302 |
| P38a | p38a MAP kinase | P38 isoform | - | 0.211 | -0.737 |
| P38b | p38b MAP kinase | P38 isoform | - | 0.133 | 0.358 |
| p38c | p38c MAP kinase | P38 isoform | - | -0.305 | -0.946 |

FC = fold change

Lines that showed a marked reduction in rab7 staining were checked for changes in bouton morphology in the double knockdown lines. All rescued MAPK flies showed a decrease in bouton size, while Thor overexpression in a foxo knockdown exhibited a reduced but non-significant change in average bouton area (Figure 4D). To validate that not all lines had smaller bouton areas, we quantified foxo;p38a double knockdown flies, and found no significant difference from foxo knockdown alone. To assess how MAPK and TGF-b impact the cytoskeleton, we performed immunofluorescence against acetylated a-Tubulin for the *foxo-RNAi/p38b-RNAi* and *foxo-RNAi/babo-RNAi* double knockdown lines. We observed that *foxo-RNAi/p38b-RNAi* flies have significantly more straight branches than the foxo-RNAi lines (Figure 4E). This was accompanied by reduced splaying along the microtubules (Figure 4F). These results support that many of the knockdowns that rescued rab7 accumulation also reduce bouton expansion, and that the b isoform of p38 is likely involved in FOXO-mediated NMJ regulation.

We next examined how motor neuron FOXO impacts MAPK activation. Using our generated ok6-foxoRNAi fly lines, we stained the A3 VLM with antibodies against the phosphorylated state of all three MAP Kinases, phosphor-Erk, phosphor-JNK, and phosphor-p38. We did not observe clear punctae for phosphor-JNK and thus could not quantify results. However, we did observe a significant increase in phospho-p38 intensity in the foxo-RNAi flies (figure 5A-B). We also observed an increase in phospo-Erk intensity in the foxo-RNAi flies, though this increase was not significant (Figure 5 C). These results show that FOXO presence in the motor neuron can suppress MAPK activation.

**Figure 5.**
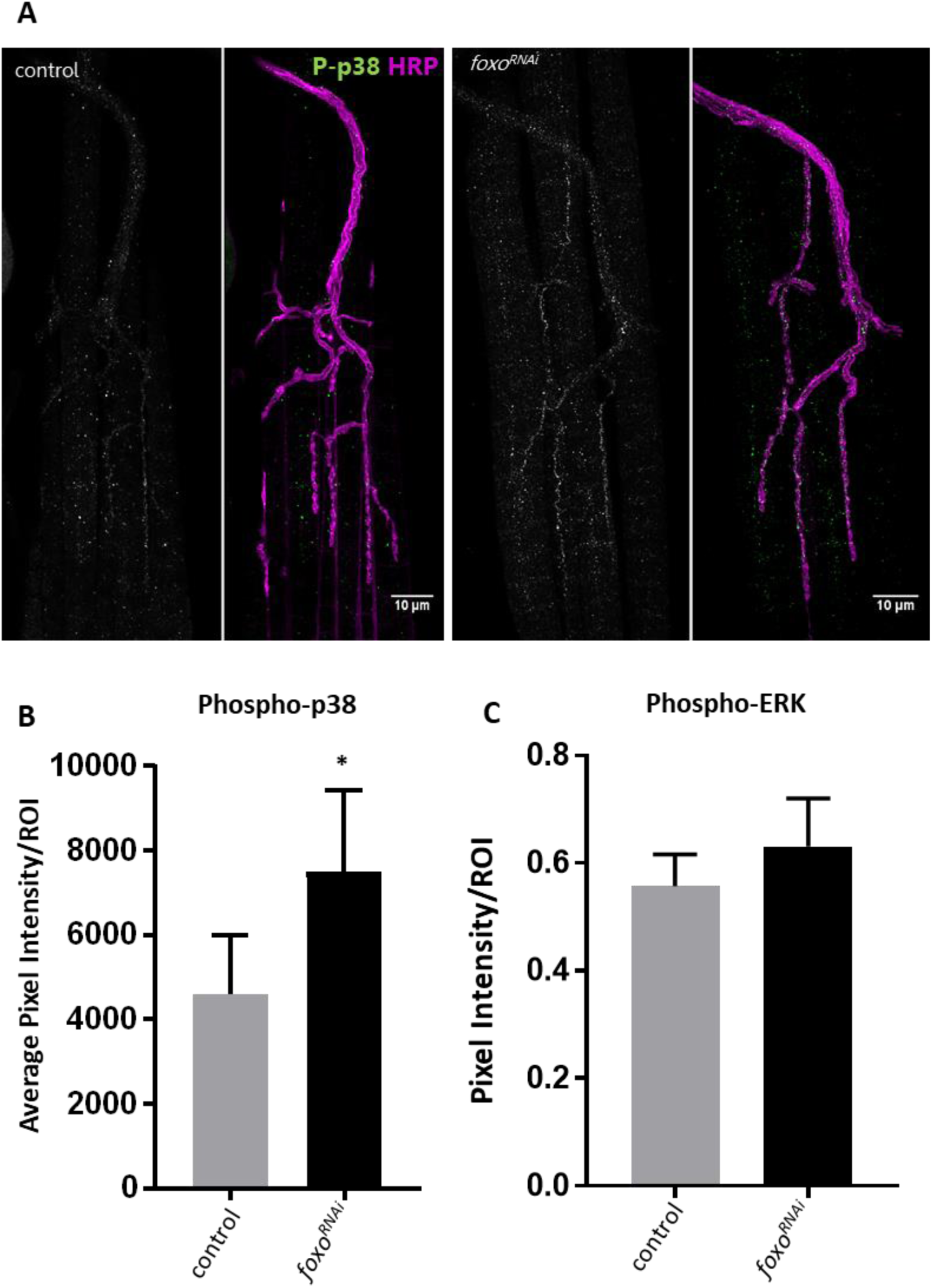
Motor neuron FOXO inhibits MAPK activation. **A.** Representative images of 1-week old fly NMJs. Colabeled with anti-phosphorylated-p38 and anti-HRP. Scale bar10μm. **B.** Relative intensity of P-p38 at anti-HRP masked areas. Intensity was normalized to anti-HRP intensity. *P<0.05. **C.** Relative intensity of P-ERK at anti-HRP masked areas. Intensity was normalized to anti-HRP intensity.

## Discussion

Foxo is known to act a regulator of synaptic plasticity and microtubule stability (Howlett et al 2008; Nechipurenko and Broiheir 2012). Although we have seen FOXO’s regulation at the NMJ during development, it is unclear the role it plays in NMJ maintenance during aging. Foxo promotes youthful characteristics in many different tissues, and has been shown to delay the onset of degenerative properties (Sahil and Brunet 2008, Desmontes and Perrimon 2012). Here we demonstrate that FOXO is a regulator of NMJ homeostasis in post developmental motor neurons. Our data suggests that foxo plays a functional role in maintaining the motor neuron during adulthood, and can promote healthy neuronal tissue when expressed later in life. We have found that foxo contributes to the morphology of synaptic contacts to muscles tissue, and can positively regulate cytoskeletal plasticity and endocytic processes in neuronal tissue. Our results also show that TGF-b and MAPK signaling function downstream of foxo to control these processes and loss of foxo may induce activation of these pathways, resulting in disruption of homeostatic mechanisms and premature cellular aging.

In this paper, we show that loss of foxo causes morphological changes to synaptic structures in young adult female drosophila. While foxo mutants have been previously shown to have larger boutons compared to controls during development, they were not reported to have any changes in branch length or number, nor were any differences reported in bouton number (Nechipurenko and Broiheir 2012, McLaughlin et al 2016). We have found this not to be the case in the adult. Our results showed that at 5 days post eclosion, foxo mutant flies had shorter branches compared to control flies. They also had a significantly larger number of branches, which ultimately resulted in a larger bouton number per muscle tissue. We hypothesize that this increase in branching may be due to a compensatory response, as foxo mutants have been shown to have reduced neurotransmission (Howlett et al 2008, Mahoney et al 2016), which may cause a need for more synaptic contacts. Additionally, *foxo* mutants are shown to have increased microtubule stability due to reduced microtubule sliding (McLaughlin et al 2016). This may inhibit growth to postsynaptic targets, resulting shorter branches. The increase in branch number is contrary to what is observed in *Drosophila* larval dendrites, where loss of FOXO reduces arborization (Sears and Broihier 2016). FOXO has also been shown to reduce neurite branching in aged *C. elegans* downstream of both JNK and Insulin signaling (Tank et al 2011). However, increases in ROS can induce overgrowth at the *Drosophila* NMJ, and FOXO is known to mitigate oxidative stress through antioxidant transcription activation (Essers et al 2004; Milton et al 2011). This increase may also be the result of ineffective pruning during metamorphosis.

Active zone number is proposed to proportionally increase with bouton size under a healthy aging model (Wagner et al 2015). Although there was no significant difference between average AZ number between controls and foxo mutants, we did observe more AZ’s per bouton in the mutants. Previous studies have shown that accumulation of Brp in axons can limit active zone number at synaptic terminals (Barber et al 2018). Foxo mutant larvae have been shown to have accumulations of Brp in the main axon (McLaughlin et al 2016), but we were unable to capture this in the adult. Based on our current data, we cannot determine whether FOXO influences active zone number. Further experimentation using Electron Microscopy of t-bars in presynaptic area could answer this question.

Aging neurons undergo cytoskeletal changes with aging in order to maintain plasticity and synaptic contact to muscle tissue (Mattson and Magnus 2006). Aging can also cause a loss of labile microtubules, while stable elements remain intact (Jones et al 2009). As previously mentioned, FOXO has been shown to regulate the axonal cytoskeleton and neuronal plasticity (Nechipuerenko, McLaughlin, Sears and Broihier 2016). During development, foxo mutant larvae have increased futch positive looping and acetylated alpha tubulin is present in the terminal boutons (Nechipurenko and Broihier 2012). Boutons were also shown to have altered Ac-Tub bundling patterns compared to controls (McLaughlin et al 2016). In our adult flies, we observed an increase in sinusoidal Ac-Tub morphologies among presynaptic branches, and this is likely a contributing factor to the truncation of established branches during aging. Dynamic stretching of axons leads to undulating distortions and mechanical failure along the axon (Tang-Schomer et al 2010). These events have been well studied in injury models, but less so in aging, where regeneration of axonal damage becomes altered (Kleele et al 2014, Hill and Coleman 2016, Geoffroy et al 2016). Taxol, a microtubule stabilizing drug, has been shown to prevent recovery from the undulating phenotype, indicating increased microtubule stability may cause the prevalence of this morphology in foxo mutants and aged controls (Tang-Schomer et al 2010). Additionally, inhibition of dynein has been shown to cause bending in microtubules (Kent et al 2016). Splayed branching in foxo mutants and motor neuron knockdown flies may be an artifact from the altered bundling pattern observed in the larvae (McLaughlin et al 2016). While the cause of this is unknown, all microtubule strands appear to be in the same phase of action, in that they are either all straight or all undulating.

Endocytosis and exocytosis play important roles in maintaining both the pre- and postsynaptic regions. Endosome to lysosome trafficking is affected in aging motor neurons and multivesicular bodies, which are prerequisites for late endosomes, accumulate (Forester et al 2010, Wagner et al 2015). Additionally, disruption of this shuttling is a known symptom of neurodegenerative diseases (Takahasi et al 2002). In our foxo knockdown flies, we found an increase in the number of late endosomes associated with the axon. This was also observed with aging. Late endosomes are upregulated for the shuttling of degraded cargo from sorting endosomes, which may indicate an increase in autophagy as a protection mechanism against harmful stress (Maiuri and Kroemer 2015; Kaur and Lakkaraju 2018). It has been shown that vesicle axonal transport declines with age, as does lysosome acidification (Forester et al 2010, Milde et al 2015, Vagnoni and Bullock 2018). Although FOXO has been shown to promote Rab7 protein expression to promote autophagy in cardiomyocytes, this was in a glucose restricted model and suggests a different mechanism than what we are observing with age (Hariharan et al 2010). However, more experimentation needs to be done to determine the underlying cause of this increase in late endosomes. Overexpression of FOXO delays late endosome increase in our adult flies. During development, a constitutively active form of FOXO caused severe microtubule destabilization, while wild-type FOXO overexpression did not have the same disruption (Nechipurenko and Boihier 2012). This may indicate a role for cytoplasmic FOXO in maintaining the integrity of motor neuron.

We first identified likely downstream targets of FOXO by examining ChIP-seq and RNA-seq data. When we conducted a genetic screen of our candidates, we found members of MAPK and TGFβ signaling to have profound rescue effects. We observed that knockdown of the activin receptor *babo* was able to rescue late endosome accumulation. TGFβ signaling has been shown to regulate axon development and Rho GTPase activity in mushroom bodies, and dictates postsynaptic density abundance at the NMJ through motor neuron anterograde signaling (Ng 2008, Kim and O’Connor 2014). TGFβ signaling has also been shown to promote aging in cardiac tissue (Chang et al 2019). We also observed that the MAP kinase p38b isoform, but not p38a or p38c, was able to rescue late endosome upregulation near the control baseline. In ALS mouse models, inhibition of p38 is able to rescue axonal retrograde trafficking (Gibbs et al 2018), and p38b is specifically shown to regulate synaptic morphology in drosophila (Klindinst et al 2013). Additionally, TGFβ has been shown to activate p38 independently of SMAD and can induce cell apoptosis through this pathway (Yu et al 2002). p38 activation has been associated with increased ROS formation and the promotion of pro-inflammatory cytokines. (Zhao et al 2018) In neuronal tissue, increased ROS and inflammation result in the activation of microglia, with chronic activation being able to induce neuronal damage and apoptosis (Dheen et al 2007; Gwak et al 2017). p38 signaling has also been shown to regulate Rab7 expression in response to interleukin cytokines (Bhattacharya et al 2006), suggesting the increase in Rab7 observed during aging and under foxo knockdown may be a response to increased inflammation.

Overexpression of the known FOXO target gene Thor had fewer rab7 punctae than foxo-RNAi alone, but did not rescue bouton morphology, suggesting this impact on late endosomes may be through a different mechanism. Additionally, some candidates such as liquid facets (*lqf*) were able to reduce the accumulation of rab7, but had severe thinning of the muscle tissue. This suggests that some genes could decrease MVB formation, but as a detriment to the health of the synapse. In our screen we also targeted *rolled* (*rl*), which is the Drosophila ortholog of ERK1/2 (Biggs et al 1994). We expected to see a decrease in late endosome accumulation, but were surprised to find there was no change. We hypothesize that this is due to the potential for two forms of ERK signaling. These two mechanisms of ERK signaling may control opposing processes of synaptic transmission. *Rolled* hypomorphs show a reduced EJP, similar to what is seen in foxo hypomorphs, suggesting reduction of *rolled* plays a part in synaptic transmission that may be independent of its role in late endosome shuttling (Wairkar et al 2009). Additionally, blockage of endosome-lysosome fusion will lead to an increase of activated ERK (Ng and Tang 2008), suggesting it may still have functional consequences downstream of FOXO activity.

In summary, we have found that loss of FOXO in the adult *Drosophila* results in morphological changes to synaptic structure that resemble aging morphologies. We also show that motor neuron FOXO is required for endocytic homeostasis, and enhanced FOXO expression during adulthood can delay aging morphologies. We find evidence to support that FOXO may act as a repressor of TGFβ and MAPK signaling, particularly p38 signaling, to maintain youthful characteristics at the NMJ. Identifying the cellular mechanisms regulated by these pathways will provide a greater understanding into how synaptic plasticity is maintained and what homeostatic processes preserve synaptic function during aging.

## Methods

### Fly stocks and husbandry

The following stocks were used: *ywR* (Rochele), *ywR;+:foxo*^*21*^, *w1118, foxoC431* (Gift from N. Liu, Chinese Academy of Sciences, Shanghai, China), *Ok6Gal4* (Gift of E McNeill, Iowa State University), *mhcGal4* (*Demontis and Perrimon* 2010, Cell). Females were collected and sorted 1-2 days after eclosion and placed in vials containing standard CSY food. Fly strains were maintained at 25°C with 12-hour light/dark cycle, and 60% humidity. Flies were transferred to fresh food every three days. Overexpression and TRiP RNAi lines were obtained from the Bloomington stock center unless otherwise specified: UAS-FOXO (BDSC_42221), FOXO-RNAi #1 (BDSC_32993), FOXO-RNAi #2 (BDSC_32427),rl-RNAi (BDSC_34855), khc-73-RNAi (BDSC_36733), lqf-RNAi (BDSC_58130), cpx-RNAi (BDSC_42017), unc-104-RNAi (BDSC_58191), UAS-Ank2GFP (Gift from Ronald Dubreuil, University of Illinois at Chicago), Bsk-RNAi (BDSC_57035), kay-RNAi (BDSC_33379), babo-RNAi (BDSC_25933), p38a-RNAi (BDSC_34744), p38b-RNAi (BDSC_29405), p38c-RNAi (BDSC_64846), grk-RNAi (BDSC_55926), bTub60D-RNAi (BDSC_64856), UAS-Thor (BDSC_9147). Fly lines w1118;ok6-Gal4 and BL32993 (FOXO-RNAi) were crossed to double balancer yw;Sp/Cyo;TM2e/TM6BTbHue (Rochele). Markers were used to select carriers. After Ok6 was combined with the foxo-RNAi, lines were backcrossed for 5 generations before use. Flies containing Ok6 in a yw background were self-crossed to produce a control strain. Line was backcrossed for 5 generations. A daughterless gene switch GAL4 driver was used to validate foxo knockdown and overexpression.

### Immunofluorescence staining

Flies were subjected to flynap (Carolina, Burlington, NC) and dissected to expose the ventral abdominal muscles (protocol from Wagner et al 2015) in Ca^2+^ free saline [128mM NaCl, 2mM KCl, 4mM MgCl_2_(H_2_O)_6_, 35.5mM sucrose, 5mM HEPES, 1mM EGTA, H_2_O, pH 7.2]. Tissue was then fixed in 4% paraformaldehyde for 20 minutes at room temperature. Tissue was washed in 1X PBST (0.1% Triton X) and blocked with 5% NGS for 1 hour at room temperature. Samples we incubated overnight at 4°C with 2ug primary antibodies in 5% NGS 1X PBST. Primary antibodies used were as follows: anti-BRP (DSHB nc82), anti-Dlg (DSHB 4F3), anti-Rab7 (DSHB), anti-rab7 (gift from Nakamura lab), anti-Acetlyated Tubulin (Sigma Aldrich T7451), anti-pERK (Cell Signaling Technologies 4370), anti-pJNK (CST 4668). Tissues we washed in 1XPBST and incubated 1.5 hours at room temperature in 1XPBST and HRP-594 or HRP-647 (Jackson ImmunoResearch Laboratories Inc, West Grove PA) at a Dilution of 1:200 with secondary antibodies (1:250) and kept in the dark at room temperature for 2 hours. Samples were washed in PBST and mounted in Prolong Diamond (Life technologies). For Fat body analysis, pelts were incubated in slowfade gold with DAPI (life technologies) at 4°C overnight prior to mounting.

### Secondary antibodies

All secondary antibodies used were from Jackson ImmunoResearch Laboratories Inc and were as follows: 488 donkey anti-mouse, 488 goat anti-rabbit, rabbit anti-HRP-594, donkey anti-HRP-647.

### Imaging analysis and Quantification

Images were captured using an epifluorescence-equipped BX51WI microscope (Olympus, Waltham, MA, USA) and Olympus FV3000 laser scanning confocal (Olympus, Waltham, MA, USA). CellSens software (Olympus, Waltham, MA, USA) was used for deconvolution of stacks. For aging metrics quantifications, maximum Z projections were generated of whole A3 muscle segment. Bouton size and active zone number were quantified in CellSense through a selected region of interest (ROI) aligned to each bouton. Measurements were performed using the “measure and count” module, providing quantification of bouton and active zone number and area. Branch number and branch length were quantified in Fiji (imageJ, Schindelin et al 2012).. Acetylated alpha-Tubulin quantification was performed via manual counting using the volume module in Fluoview software. Rab7 quantification was performed in CellSense using a circle we generated a circular ROI (25.75 µm2) to quantify number of punctae. Z-slices quantified were limited to those containing HRP staining at selected regions to avoid non-neuronal associated punctae. ROIs were checked prior to quantification to ensure muscle associated punctae were removed. Intensities for pp38, pERK, and pJNK were conducted using Fiji (NIH) by measuring the sum fluorescence intensity in ROIs defined by HRP immunoreactivity. HRP fluorescence intensity was measured to ensure no differences between control and mutant NMJs. All images were generated using Fiji.

### Quantitative real-time PCR

3-day post eclosion flies were placed on food containing less than 0.1% ethanol and 200uM Mifepristone (Cayman Chemical) or no other additive for 3 days. Whole bodies of female flies were homogenized in TRIzol (Thermo Fisher). RNA was extracted followi ng company procedure. cDNA was generated using qScript (Quanta) and diluted to a 10uM working solution. Quantitative PCR was run on purified samples (QuantStudio). Cybergreen (Life Technologies, CA, USA) was used for chemical detection. The primers used Enrichment was determined based on the double-delta CT value. *Thor* (sense, 5’ –CCAGGAAGGTTGTCATCTCGG-3’, and antisense 5’ – CTCGTAGATAAGTTTGGTGCCTCC-3’), *Rpl32* (sense 5’-AAGAAGCGCACCAAGCACTTCATC-3’, and antisense 5’ – TCTGTTGTCGATACCCTTGGGCTT-3’).

### Video capturing

Stacks of the entire A3 muscle region were acquired with Olympus FV3000 laser scanning confocal in the Fluoview software (Olympus, Waltham, MA, USA). Videos were recorded of 3D renderings using the movie module under the volume setting.

### Venn Diagrams

Venn comparisons were performed using the Bioinformatics and Evolutionary Genomics Venn calculator at http://bioinformatics.psb.ugent.be/webtools/Venn/

### Statistical analysis

GraphPad Prism (GraphPad Software) was used for statistical analysis. To compare the mean value of treatment groups versus that of control, either student t-test or one-way ANOVA was performed using Tukey multiple comparison. All genetic screen lines were compared to the generated foxo-RNAi line using a student t-test. The metrics during aging were analyzed by two-way ANOVA, including Tukey multiple comparisons test. Before analysis, outliers were identified using Robust regression and Outlier removal (ROUT) method (Q = 1%).

## Acknowledgments

We thank Bloomington Drosophila Stock Center (BDSC) and Norbert Perrimon and others at the Drosophila RNAi Screening center (DRSC) for providing fly lines. We also thank Nan Liu and Xiaofen Wu (Chinese Academy of Sciences, Shanghai, China) for fly lines and RNA-seq data, and Elizabeth McNeill (Iowa State University, Ames, IA) and Ronald Dubreuil (University of Illinois at Chicago, Chicago, IL) for fly lines used in this study. We thank Akira Nakamura (RIKEN Center for Developmental Biology, Kobe, Japan) for antibodies used in this study. The authors acknowledge the assistance of rotation student Sean McLaughin. This work was supported by NIH/NIA R00 AG048016 to HB, AFAR Research Grants for Junior Faculty to HB.

