## Supplemental figures for "FOXO regulates neuromuscular junction homeostasis during *Drosophila* aging"

**Figure S1. Raw counts for NMJ metrics. A.** bouton area **B.** branch length **C.** active zone number per bouton **D.** bouton area for foxo$^{C431}$ **E.** Branch length for foxo$^{C431}$. *P< 0.05, ** P<0.01, ***P>0.001, ****P< 0.0001, n.s. not significant.

**Figure S2. A.** Representative images of anti-Ac-Tub staining. **B.** Average intensity of Ac-Tub at synaptic terminal regions. **C.** Confocal Video capturing of 3D NMJ structure. Colabeled with anti-Ac-Tub and anti HRP.

**Figure S3. A.** Rab7 quantification at 1 week for ok6>ywR, ok6>foxo-RNAi #1, ok6> foxo-RNAi #2. **B.** Thor relative expression for whole body tissue. Daughterless-Gal4 activated with 200mM of Mifepristone.

**Figure S4. Representative confocal images from genetic screening. A.** anti-rab7 and anti-HRP immunofluorescence for ok6;foxo$^{RNAi}$> p38b$^{RNAi}$, ok6;foxo$^{RNAi}$> rl$^{RNAi}$, ok6;foxo$^{RNAi}$> p38a$^{RNAi}$. **B.** anti-Ac-Tub and anti-HRP immunofluorescence for ok6;foxo$^{RNAi}$> p38b$^{RNAi}$, ok6;foxo$^{RNAi}$> rl$^{RNAi}$, ok6;foxo$^{RNAi}$> p38a$^{RNAi}$.

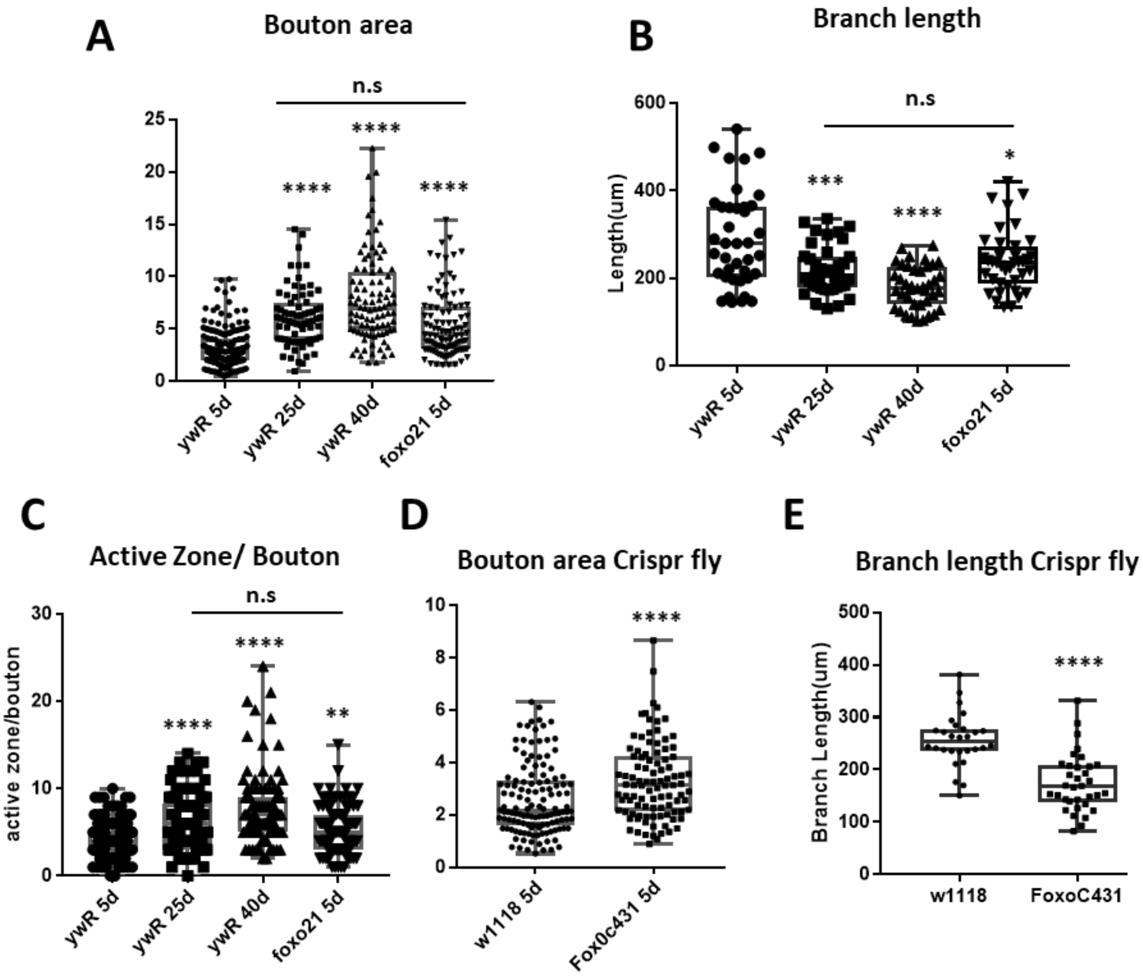
A
Bouton area
n.s
****
****
****
0
5
10
15
20
25
ywR 5d
ywR 25d
ywR 40d
foxo21 5d
B
Branch length
n.s
***
****
*
Length(um)
0
200
400
600
ywR 5d
ywR 25d
ywR 40d
foxo21 5d
C
Active Zone/ Bouton
n.s
****
****
**
active zone/bouton
0
10
20
30
ywR 5d
ywR 25d
ywR 40d
foxo21 5d
D
Bouton area Crispr fly
****
0
2
4
6
8
10
w1118 5d
Fox0c431 5d
E
Branch length Crispr fly
****
Branch Length(um)
0
100
200
300
400
500
w1118
FoxoC431

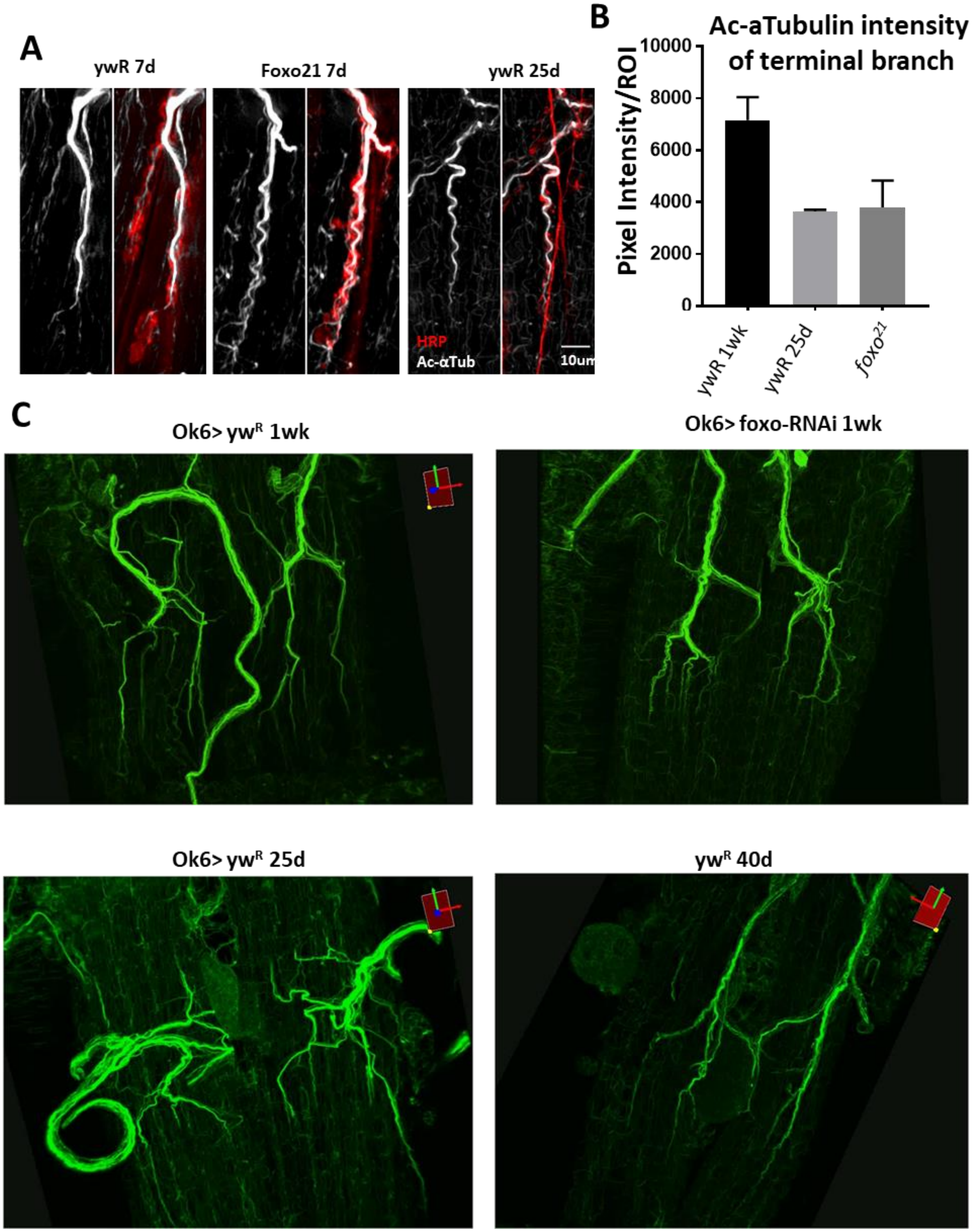
A
ywR 7d
Foxo21 7d
ywR 25d
HRP
Ac-αTub
10um
B
Ac-aTubulin intensity of terminal branch
Pixel Intensity/ROI
10000
8000
6000
4000
2000
0
ywR 1wk
ywR 25d
$foxo^{21}$
C
Ok6> yw$^{R}$ 1wk
Ok6> foxo-RNAi 1wk
Ok6> yw$^{R}$ 25d
yw$^{R}$ 40d

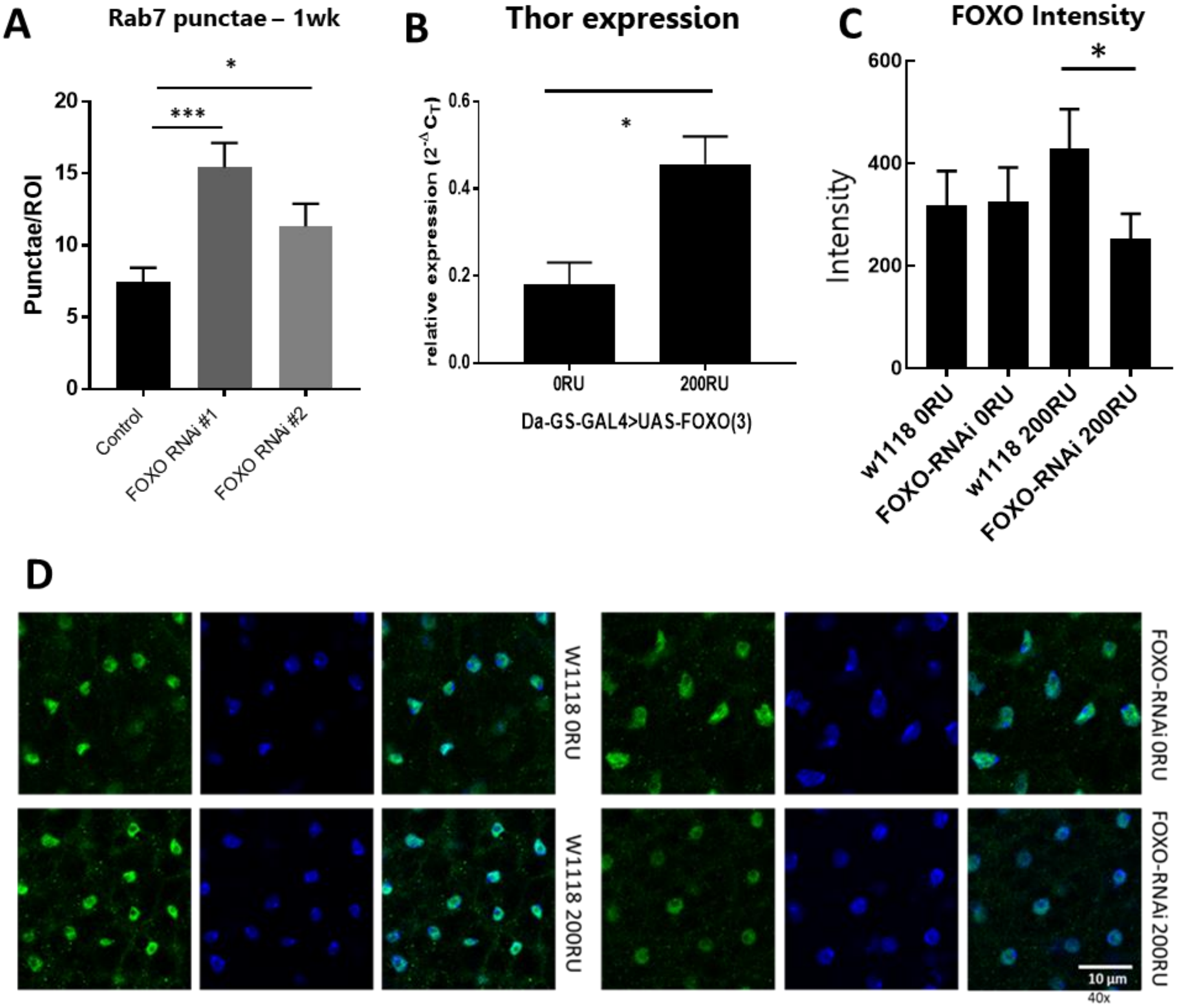

A
Rab7 punctae – 1wk
Punctae/ROI
20
15
10
5
0
***
*
Control
FOXO RNAi #1
FOXO RNAi #2
B
Thor expression
relative expression (2-ΔCT)
0.6
0.4
0.2
0.0
*
0RU
200RU
Da-GS-GAL4>UAS-FOXO(3)
C
FOXO Intensity
Intensity
600
400
200
0
*
w1118 0RU
FOXO-RNAi 0RU
w1118 200RU
FOXO-RNAi 200RU
D
W1118 0RU
W1118 200RU
FOXO-RNAi 0RU
FOXO-RNAi 200RU
10 µm
40x

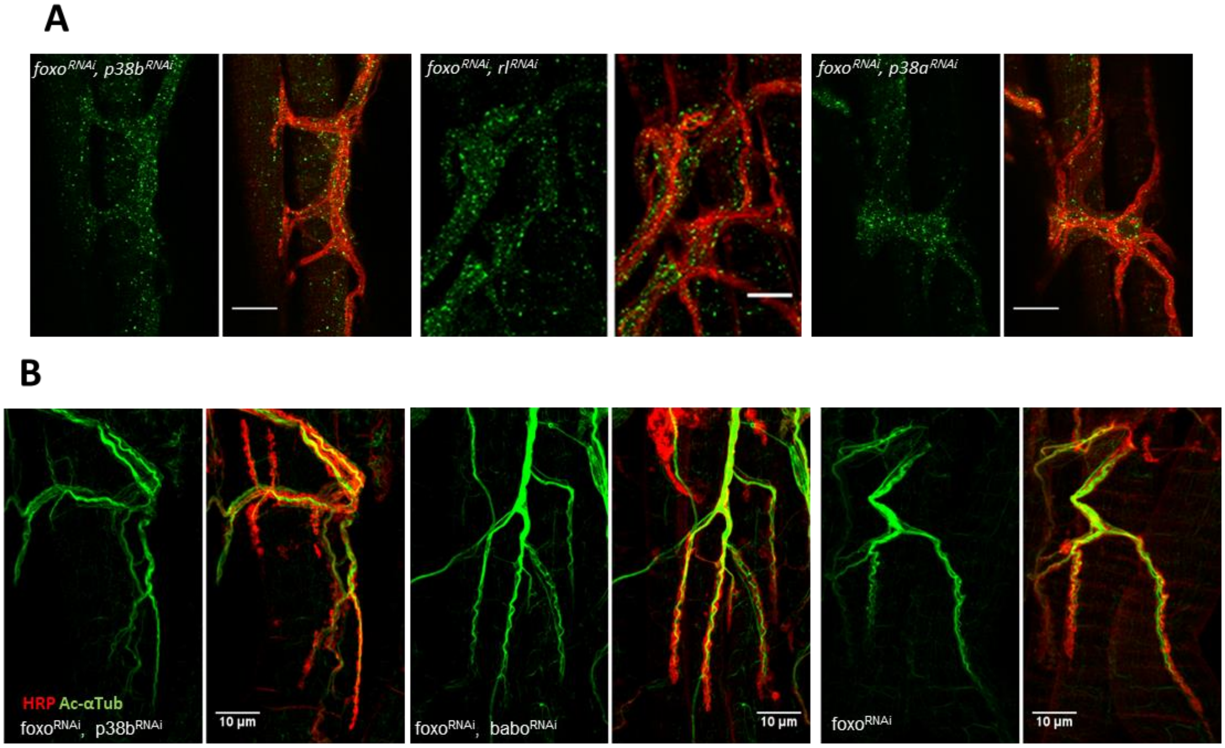
A
foxo$^{RNAi}$, p38b$^{RNAi}$
foxo$^{RNAi}$, rl$^{RNAi}$
foxo$^{RNAi}$, p38a$^{RNAi}$
B
HRP Ac-αTub
foxo$^{RNAi}$, p38b$^{RNAi}$
10 μm
foxo$^{RNAi}$, babo$^{RNAi}$
10 μm
foxo$^{RNAi}$
10 μm
